# The relationship between white matter architecture and language lateralisation in the healthy brain

**DOI:** 10.1101/2024.01.12.572159

**Authors:** Ieva Andrulyte, Christophe De Bezenac, Francesca Branzi, Stephanie J Forkel, Peter N. Taylor, Simon S. Keller

**Affiliations:** Department of Pharmacology and Therapeutics, Institute of Systems, Molecular and Integrative Biology, University of Liverpool, Liverpool, UK; The Walton Centre NHS Foundation Trust, Liverpool, UK; Department of Psychological Sciences, Institute of Population Health, University of Liverpool, Liverpool, UK; Donders Institute for Brain Cognition Behaviour, Radboud University, Nijmegen, the Netherlands; Brain Connectivity and Behaviour Laboratory, Sorbonne Universities, Paris, France; Centre for Neuroimaging Sciences, Department of Neuroimaging, Institute of Psychiatry, Psychology and Neuroscience, King’s College London, London, UK; CNNP Lab (www.cnnp-lab.com), Interdisciplinary Computing and Complex BioSystems Group, School of Computing Science, Newcastle University, UK; Institute of Neurology, Queen Square, UCL, London UK

**Keywords:** white matter asymmetries, language lateralisation, tractography, connectometry, hemispheric asymmetries, MRI, corpus callosum, forceps minor

## Abstract

Interhemispheric anatomical asymmetries have long been thought to be related to language lateralisation. Previous studies have explored whether asymmetries in the diffusion characteristics of white matter language tracts are consistent with language lateralisation. These studies, typically with smaller cohorts, yielded mixed results. This study investigated whether connectomic analysis of quantitative anisotropy (QA) and shape features of white matter tracts across the whole brain are associated with language lateralisation. We analysed 1040 healthy individuals from the Human Connectome Project database. Hemispheric language dominance for each participant was quantified using a laterality quotient (LQ) derived from fMRI activation in regions of interest (ROIs) associated with a language comprehension task compared against a math task. A linear regression model was used to examine the relationship between structural asymmetry and functional lateralisation. Connectometry revealed that LQs were significantly negatively correlated with QA of corpus callosum tracts, including forceps minor, body, tapetum, and forceps major, indicating that reduced language dominance (more bilateral language representation) is associated with increased QA in these regions. The QA of the left arcuate fasciculus, cingulum, and right cerebellar tracts was positively associated with LQ, suggesting that stronger structural asymmetry in these tracts may identify left language dominance. Language lateralisation was not significantly associated with the shape metrics (including length, span, curl, elongation, diameter, volume, and surface area) of all white matter tracts. These results suggest that diffusion measures of microstructural architecture, and not the geometric features of reconstructed white matter tracts, are associated with lateralisation of language comprehension functions. People with increased dependence on both cerebral hemispheres for language processing may have more developed commissural fibres, which may support more efficient interhemispheric communication.

## Introduction

It has long been hypothesised that neuroanatomical asymmetries of regions that support language may be associated with functional lateralisation of language in the human brain (Güntürkün et al., 2020). Historical evidence dates to post-mortem studies of acquired language disorders in the 19th century, which identified the involvement of the left frontal, temporal, and parietal lobes in language function (Broca, 1861; Dax, 1863; Wernicke, 1874). Some cytoarchitectonic, post-mortem, and quantitative magnetic resonance imaging (MRI) studies have reported leftward asymmetry of language cortical regions in individuals with unknown hemispheric language dominance (HLD) (Amunts & Zilles, 2006; Uylings et al., 2006; Foundas et al., 1998; Keller et al., 2007; Galaburda et al., 1978; Geschwind & Levitsky, 1968), whereas others have reported no asymmetry (Good et al., 2001; Keller et al., 2009; Luders et al., 2004; Tomaiuolo et al., 1999; Wada et al., 1975; Watkins et al., 2001).

Studies examining the association between cerebral asymmetry and HLD are equally mixed. Associations between localised cerebral asymmetries and HLD have been noted in some studies of Wada-tested patients with epilepsy (Dorsaint-Pierre et al., 2006; Foundas et al., 1996, 2002; Keller et al., 2018) and healthy controls who underwent structural and functional MRI (Josse et al., 2009; Keller et al., 2011). However, other studies reported no association between HLD and structural hemispheric asymmetry in classical language areas (Chiarello et al. 2013; Greve et al. 2013).

Several structural MRI and diffusion tensor imaging (DTI) studies have examined the relationship between HLD and white matter volume and have reported that leftward language lateralisation is associated with a greater volume of the arcuate fasciculus (Propper et al., 2010) and the number of tracts in the corpus callosum (Timocin et al., 2020). More recently, HLD has been investigated using microstructural DTI properties, such as fractional anisotropy (FA), in the context of the presurgical evaluation of epilepsy and tumours (Tantillo et al., 2016; Barba et al., 2020). Some studies have reported relationships between language lateralisation and interhemispheric asymmetry of diffusion characteristics or the size of the corpus callosum (Tantillo et al., 2016), while others did not (Barba et al., 2020). A fundamental issue in presurgical patient studies is the increased prevalence of functional and structural reorganisation of language networks and abnormal language lateralisation (Yogarajah et al., 2010). There are limited functional MRI (fMRI)-DTI studies on language lateralisation in healthy individuals. Most existing studies have used similar tractography methods, and some have reported significant correlations between the fMRI-determined left HLD and FA of the left arcuate fasciculus (James et al., 2015; Perlaki et al., 2013; Powell et al., 2006; Silva & Citterio, 2017) and corpus callosum (Häberling et al., 2011). However, other studies have not reported relationships between the side or extent of HLD and conventional diffusion-based tract characteristics (Karpychev et al., 2022; Vernooij et al., 2007), as well as more sophisticated diffusion MRI measures, such as fibre density cross-section (Verhelst et al., 2021). Inconsistencies between studies may be due to methodological differences, including differences in tractography approaches, study designs, patient characteristics, and sample sizes.

Most diffusion studies investigating the white matter basis of HLD used conventional tractography approaches to determine inter-hemispheric asymmetries of the overall tract size or diffusion metrics averaged over entire tracts (Perlaki et al., 2013; Vernooij et al., 2007). In the present study, we adopted two complementary approaches that could potentially overcome some of the methodological shortcomings of previous tractography studies. First, we employed a connectometry approach based on an automated fibre-tracking algorithm, which uses permutation testing to identify group differences along white matter tracts. This whole-brain approach employs correlational tractography to identify specific subcomponents of white matter tracts that exhibit anisotropy correlated with a predefined variable of interest with superior sensitivity and specificity compared to traditional voxel-based analyses (Yeh et al., 2016). Connectometry has recently been used to uncover structural disparities between bilingual and non-bilingual individuals (Rahmani et al., 2017) and to identify structural pathways linked to enhanced language capabilities in individuals with aphasia (Hula et al., 2020; Dresang et al., 2021) and preterm-born children (Barnes-Davis et al., 2020, 2022).

However, it has not been used to determine white matter structural correlates of HLD. Second, we employed shape analysis to investigate the geometrical characteristics of white matter tract bundles that comprise the integral components of language networks. This approach captures fundamental shape characteristics, such as volume and surface area, and encompasses advanced morphological properties including white matter bundle curl, elongation, length, span, and diameter (Yeh, 2020). Grey matter studies have highlighted the significance of examining the rich morphology in cortical areas, particularly the developmental patterns of sulci and gyri, revealing that leftward asymmetries in the left inferior frontal gyrus may arise from additional branches, such as the diagonal sulcus (Keller et al., 2007). However, little research has been conducted on the relationship between HLD and white matter morphometry. Previous studies have already demonstrated, through the utilization of virtual dissections (Catani et al., 2007) and shape analysis employing tractography algorithms (Yeh et al., 2020), that the leftward asymmetry of language-associated white matter tracts exists, notably of the arcuate fasciculus and uncinate fasciculus. However, the association between these asymmetries and HLD remains unclear.

Our study had two main objectives. First, we conducted diffusion connectometry analysis in a large cohort of healthy individuals who underwent assessment of language function using fMRI to determine whether localised parts of white matter tracts are related to HLD across the entire brain. This enabled us to determine whether regional QA is related to a lateralised function without being limited to averaging diffusion scalar metrics over the entire length of each tract. The second objective was to explore whether interhemispheric shape asymmetries of the white matter tracts are related to language lateralisation in the same individuals.

## Methods

### Study data and participants

All data were acquired from the Human Connectome Project (HCP) (http://www.humanconnectome.org/) open-access data initiative offering high-quality anatomical and functional MRI of the human brain. We used the HCP Young Adults (HCP-YA 1200 Subjects) data release as it contains a large sample of healthy adults for whom both language task fMRI and diffusion MRI sequences were acquired. The dataset comprised 1200 healthy adults, aged 22-35 years. Each participant underwent an identical imaging protocol acquired on the same MRI scanner. Individuals with neuropsychiatric or neurologic disorders, diabetes, high blood pressure, premature birth, and severe symptoms associated with substance use were excluded from data collection (Van Essen et al., 2013). The present study focused on language fMRI and diffusion MRI data only. Individuals were only selected for inclusion if they had fMRI data available for the language story task (see below) and had corresponding 3T diffusion MRI data. This resulted in a sample size of 1040 participants (562 females), with a mean age of 28.74 (SD = 3.69) years. According to the Edinburgh Handedness Inventory (Oldfield, 1971), 962 (92%) participants preferred their right hand, scoring at least 10 on a scale of −100 (left) to 100 (right). Eighty-five participants preferred left, scoring below −10, and two were ambidextrous, scoring zero.

### Data acquisition

HCP data were acquired on a Siemens 3T Skyra system, with 32-channel (SC72) head coil. Task fMRI data were collected using gradient-echo echo-planar imaging (EPI) with an isotropic resolution of 2.0 mm (TR = 720ms, TE = 33.1ms, matrix = 104×90, 72 slices, flip angle = 52°, BW = 2290 Hz/Px, FOV = 208 x 180 mm, 72 slices, multiband accelerator factor = 8) (Marcus et al., 2013). The HCP dMRI data were acquired using three shells (b=1000, 2000 and 30009 s/mm2) with 90 diffusion gradient directions and five b_0_ volumes with RL phase encoding direction (TE = 89.5ms, TR = 5520ms, flip angles = 78/160°, isotropic voxel size = 1.25 mm3, multiband factor = 3) (Sotiropoulos et al., 2013).

### Language paradigm

The language comprehension task used in Human Connectome Project was designed by Binder and colleagues (2011). The task consists of two 3.8-minute runs. Each run has four blocks of a story task alternating with four blocks of a math task. The story and math tasks are matched in terms of length, word and phoneme rate, speaking style, and prosodic features. While the story blocks present subjects with 5-9 auditory sentences, followed by questions about the content of the story, the math task requires participants to perform arithmetic operations followed by equals and two choices. This contrast allowed us to cancel out the regions that are jointly activated in both tasks, isolating the regions involved in narrative processing including semantic and syntactic integration, theory of mind, and inference processing.

### fMRI preprocessing and analysis

The preprocessed task fMRI data were retrieved from HCP database (https://db.humanconnectome.org). The HCP pre-processing included *fMRIVolume* and *fMRISurface* pipelines, which were primarily built using tools from FSL (Jenkinson et al., 2012; http://www.fmrib.ox.ac.uk/fsl), Freesurfer (Fischl, 2012) and the HCP Workbench (Marcus et al., 2013). Details of the pre-processing steps have been described previously (Glasser et al., 2013). The goal of the first *fMRIVolume* pipeline was to generate 4D whole-brain timeseries. This was accomplished by (1) removing spatial distortions by gradient non-linearity distortion correction, (2) realigning volumes using rigid-body motion correction using a single-band reference image as the target, and (3) estimating (using FSL toolbox “topup”) and correcting field map-based EPI distortions. The resulting EPI data was (4) registered to T1-weighted scan, then (5) non-linearly (FNIRT) into Montreal Neurological Institute (MNI) space, and (6) blood-oxygen-level−dependent (BOLD) signal intensity was normalized by the average.

The goal of *fMRISurface* pipeline was to transform the resulting 4D timeseries to Connectivity Informatics Technology Initiative (CIFTI) grayordinate space, encompassing cortical, subcortical, and cerebellar grey matter collectively (Pham et al, 2022). This was accomplished by mapping fMRI data within cortical grey matter ribbon onto the native cortical surface, registering it into CIFTI grayordinate space (surface representation with 32,492 vertices on each hemisphere), and mapping the set of subcortical grey matter voxels from each subcortical parcel in each individual to a standard set of voxels in each atlas parcel, resulting in 2mm average surface vertex and subcortical volume voxel spacing. Finally, grayordinate space data was smoothed using Gaussian kernel.

We used a fully processed task-based STORY-MATH fMRI activation Contrast Of Parameter Estimates (COPE) map, which was generated by FSL FEAT and is readily available on https://db.humanconnectome.org as part of the “S1200 Subjects” dataset. Considering the spatial heterogeneity of the individual brain scans, the MSM-All (Multimodal Surface Matching) registered dataset was used, which uses information on areal features derived from the resting state network, myelin maps, and alignment of folding. The motivation for using MSM-All over MSM-Sulc (cortical folding-based registration) came from previous studies that demonstrated a weaker correlation between sulcal depth and local curvature with regions responsible for higher cognitive functions, including Broca’s area (Fischl et al., 2008; Van Essen, 2005), compared to the MSM-All registration, which showed improved cross-subject alignment of independent task fMRI datasets (Robinson et al., 2018).

### Laterality Quotient

Grayordinates localised regions of interest (ROIs) on the “inflated” brain surface (Van Essen & Glasser, 2016). A laterality quotient (LQ) was calculated to assess HLD for each participant’s task fMRI activation using the CIFTI toolbox in MATLAB in ROIs associated with language comprehension. We decided to use anterior language regions of interest (ROIs), specifically Brodmann areas 44 and 45, due to their demonstrated higher reliability in determining language dominance in semantic tasks (Seghier et al., 2008; Seghier, 2008).

Subsequently, the selection of these ROIs was further substantiated upon closer examination of individual activation maps. In these maps, it became apparent that temporal regions exhibited predominantly bilateral activation in cases of both left and right hemisphere dominance. However, a notable asymmetry emerged in frontal activation, particularly when focusing on the specific semantic comprehension paradigm. This phenomenon aligns with previous research by Sabbah and colleagues (2003), wherein the activation of frontal lobes during semantic tasks was the sole activation consistently correlated with the results of the WADA test. Consequently, this serves as additional validation for our deliberate choice of ROI (Figure 1).

**Figure 1.**
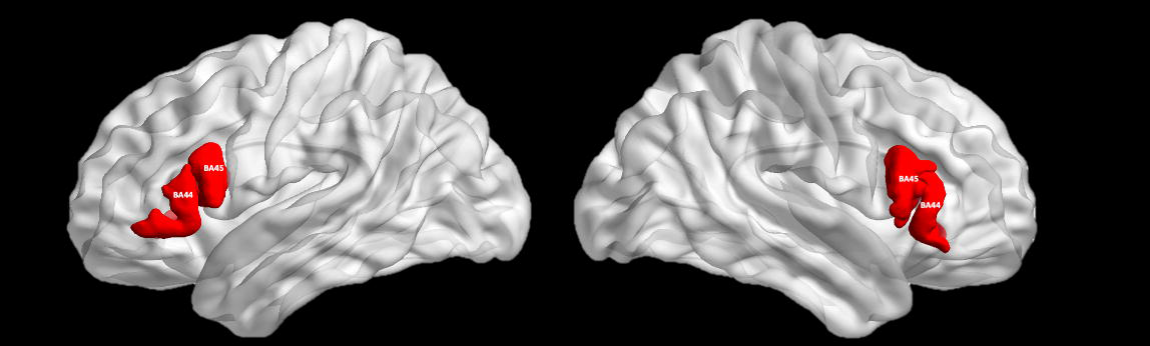
Inferior frontal areas (BA44, BA45) were selected as regions of interest to calculate LQ. The z-values of the fMRI activations on the left side were subtracted from the z-values of the fMRI activations on the left side and then divided by the sum of the z-values on both hemispheres. The LQ value is expressed as a percentage, ranging from −1 to 1, where > 0.1 refers to left-lateralized, < −0.1 to right-lateralized, and the values between −0.1 and 0.1 to bilateral individuals.

To work with CIFTI files, we generated dscalar files for each ROI using wb_command, imported them into MATLAB, and extracted z-values from ROIs using the CIFTI toolbox. LIs were then measured for each participant’s active greyordinates (Z > 1.96, corresponding to two-tailed uncorrected P < 0.05) using the equation below:

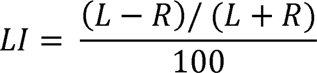

Where L represents the number of active voxels in the left, and R in the right hemisphere. Given that the LQ ranges from −1 to +1, values above +0.1 indicate left language dominance (LLD), values below −0.1 indicate right language dominance (RLD), and values between −0.1 and +0.1 indicate bilateral language representation (BLR) (Yuan et al, 2006).

### Diffusion processing

Diffusion data was downloaded from the HCP S1200 Young Adult Data Release and preprocessed using the HCP Diffusion preprocessing pipeline using FMRIB diffusion toolbox in FSL. Briefly, the pipeline included b_0_ image intensity normalisation, removing EPI susceptibility-induced field distortions with FSL’s “topup” algorithm (Andersson et al, 2016), correcting for eddy current distortions, head movements, and gradient nonlinearities (Glasser et al., 2013). Quality control of the preprocessed diffusion MRI data was performed using DSI studio software (http://dsi-studio.labsolver.org). An automatic quality control routine then checked the b-table to ensure its accuracy (Schilling et al., 2019). The diffusion data were co-registered in MNI space using q-space diffeomorphic reconstruction (Yeh & Tseng, 2011) to obtain the spin distribution function (SDF) with a recommended length ratio of 1.25, as specified in the original study (Yeh et al., 2010) (Figure 2).

**Figure 2.**
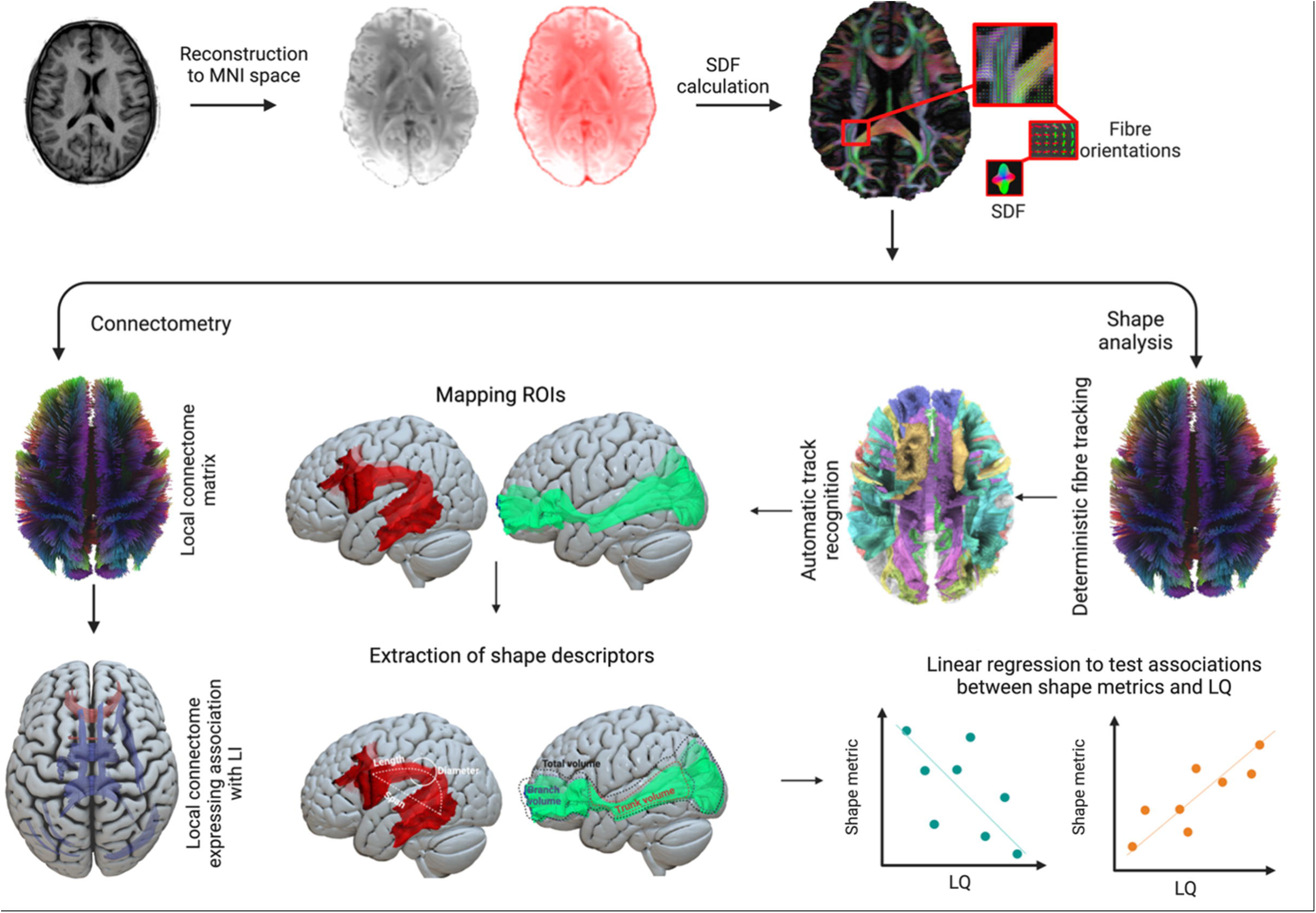
Flowchart of the methods pipeline. The pre-processed diffusion MRI data was reconstructed in an MNI space. The outputs of the reconstruction and SDFs were calculated to obtain the fibre orientations using DSI studio. Then, two different approaches were used to examine the white matter tracts associated with language laterality. The connectometry approach involved obtaining a local connectome matrix and finding out its association with LQ. Shape analysis involved the recognition of the WM tracts using HCP atlas and mapping eleven WM fibre bundles important for language function. The measures of key shape features, such as curl and volume, were extracted and linear regression analyses were used to look at the associations between shape metrics and LQ.

### Connectometry analysis

We applied group connectometry analysis using DSI Studio as described in previous applications (Barnes-Davis et al., 2020, 2022; Dresang et al., 2021; Rahmani et al., 2017) to study the relationship between regional white matter quantitative anisotropy (QA) and language lateralisation measures derived from LQs (Figure 2). The connectometry approach derives the QA measure from the SDF in each fibre orientation, which defines the number of anisotropic spins along that direction in each streamline (Yeh et al., 2010, 2013). The anisotropy in each section of a white matter tract is then correlated with the study variable (Yeh et al., 2016). Unlike a voxel-based FA metric, which attributes identical anisotropy values to all fibre orientations within a voxel, QA demonstrates a discerning capability by identifying specific axonal orientations in each peak orientation of the SDF. This selectivity effectively distinguishes less prominent peaks and mitigates their influence, thereby reducing the risk of false tracking and further elevating spatial resolution (Yeh et al, 2013).

Our connectometry analyses were conducted in three stages: the first stage of the connectometry analysis included only participants with LLD and BLR; the second stage included participants with RLD and BLR; and the third stage involved participants with both LLD and RLD. We chose not to perform the regression analyses on all groups simultaneously to facilitate interpretation and to capture the distinct effects associated with different degrees of laterality. The linear effect of handedness, sex, and age was mitigated using a partial linear correlation. A nonparametric Spearman partial correlation was used to derive the continuous segments correlating with a LQ (Yeh et al., 2016). Each reconstructed white matter tract within a voxel was tracked to extract a QA map in the native diffusion space for each participant (Yeh et al., 2013). A T-score threshold was assigned to the highest level of three to reduce the possibility of false positive results (Ashraf-Ganjouei et al., 2019). The tracks were filtered by topology-informed pruning with sixteen iterations to remove implausible spurious connections (Yeh et al., 2019). A false discovery rate (FDR) correction for multiple comparisons was employed with a threshold of 0.05 to select tracks showing significant associations between LQ and QA with a default length threshold of 15 voxels (Barnes-Davis et al., 2020). To estimate the false discovery rate, 4000 randomized permutations were applied to the group label to obtain the null distribution of the track length. After the correlational results were obtained, additional categorical analyses were performed at the group level (LLD/RLD, LLD/BLR, RLD/BLR).

To better reflect the true anatomy of the streamlines, the substantial tracts were displayed in their entirety as a complete bundle. Additionally, raw images were made available to showcase only the most important parts of the tract.

### Shape analysis

The SDF maps generated from the connectometry analysis were used for tract shape analysis. Each participant’s SDF map was normalised to MNI space using nonlinear registration, and automatic fibre tractography was performed using a deterministic fibre tracking algorithm utilising DSI studio software (Yeh, 2020). Eleven white matter tract bundles that are part of language comprehension networks (Forkel et al, 2022; Friederici et al., 2007; Harvey et al., 2013; Ivanova et al., 2021; Rollans & Cummine, 2018; Shin et al., 2019; Zhong et al., 2022;) were then automatically tracked using the HCP1065 tractography atlas (Figure 3). These include the arcuate fasciculus (AF), corpus callosum body, corpus callosum forceps major (splenium), corpus callosum forceps minor (genu), inferior fronto-occipital fasciculus (IFOF), frontal aslant tract (FAT), inferior longitudinal fasciculus (ILF), the three branches of the superior longitudinal fasciculus (dorsal SLF1, middle SLF2 and ventral SLF3), and the uncinate fasciculus. All white matter bundles were independently tracked within the left and right hemispheres, while the corpus callosum bundles were tracked as a whole. The diffusion sampling length ratio was set at 1.25 and the output resolution was resampled to 2 mm isotropic. To remove false connections, topology-informed pruning was applied with 32 iterations (Yeh et al., 2019). Finally, after identifying all white matter tracts of interest, several shape metrics including tract length, span, curl, elongation, diameter, volume, and surface area were extracted (Figure 4).

**Figure 3.**
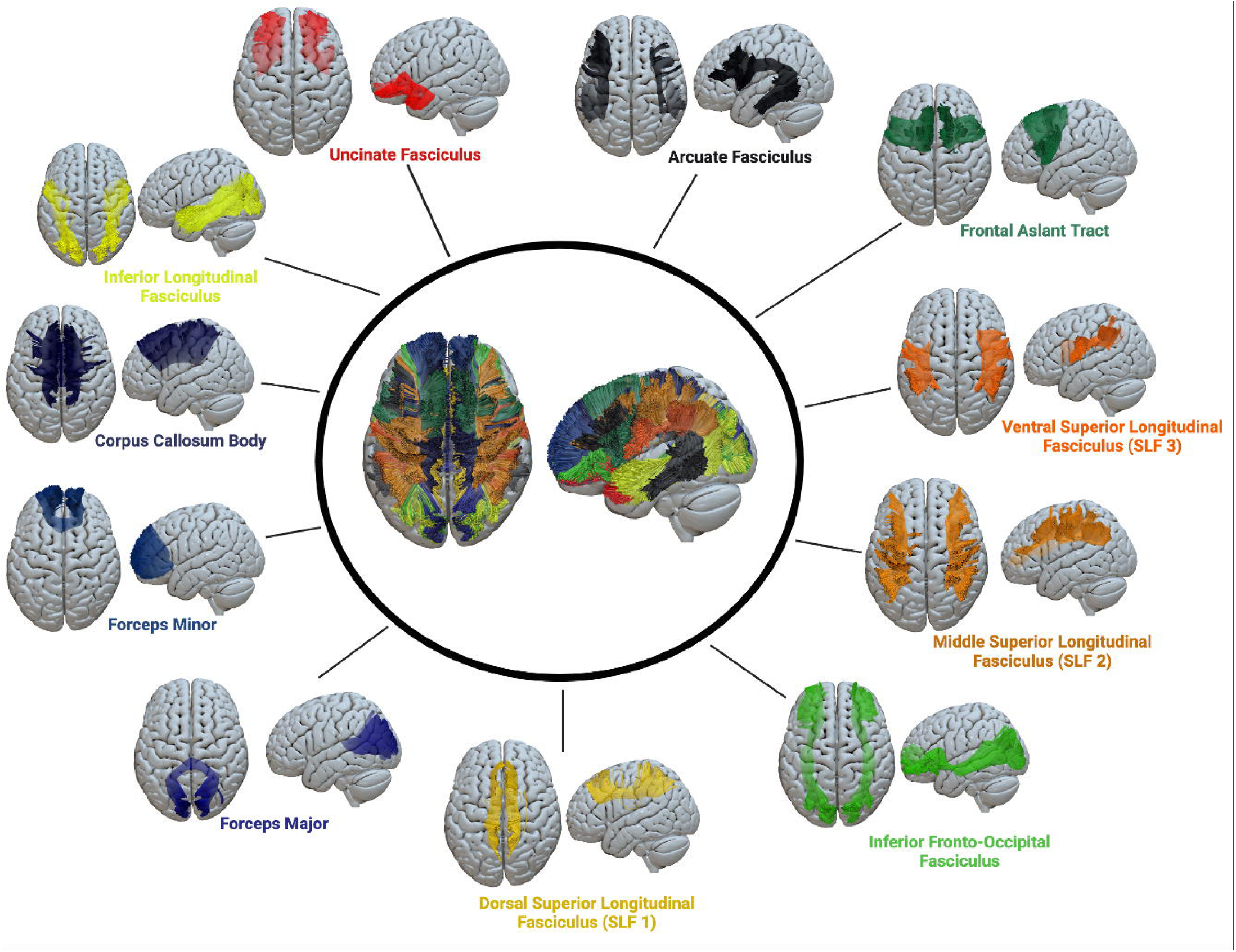
Eleven white matter tracts were reconstructed for shape analysis based on the HCP842 atlas computed on 1065 healthy people (Yeh et al, 2018).

**Figure 4.**
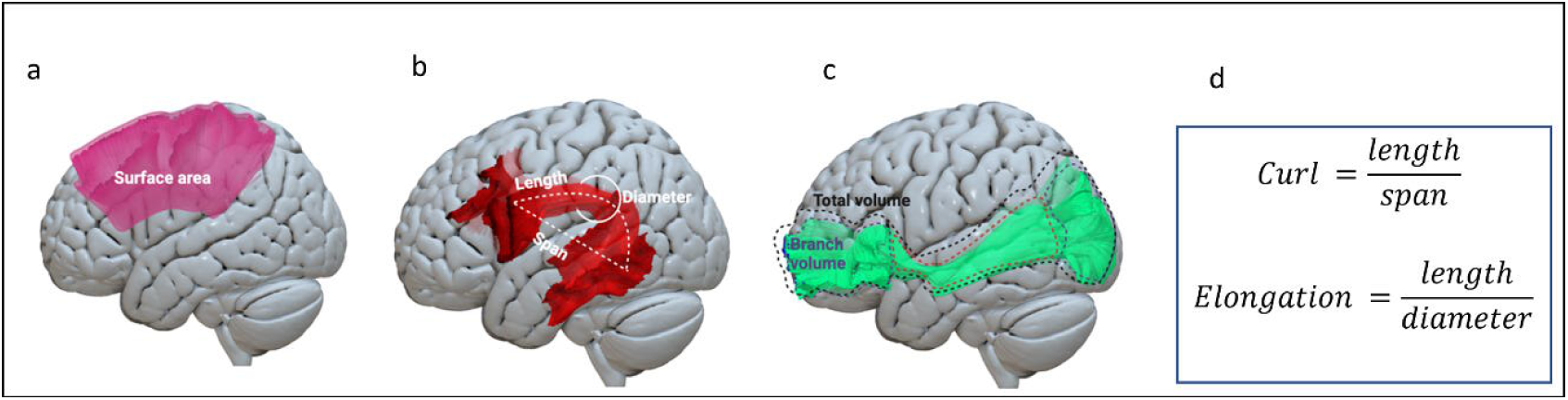
Schematic illustration of the shape analysis of the white matter tracts. (a) The area metrics used in the included surface area (mm). (b) The length metrics used in the study included mean tract length (mm) as well as span bundle (mm) and diameter (mm) of the bundle. (c) The volume metrics used in the study included branch volume (mm^3^), trunk volume (mm^3^), and total bundle volume (mm^3^). (d) The shape metrics used in the study included curl and elongation.

To investigate the relationship between the shape of the white matter tracts and functional language laterality, linear regression analysis was performed using the same groups and covariates as in connectometry analyses. We also conducted an analysis of variance (ANOVA) to assess the statistical significance of differences among different laterality groups (LLD/RLD/BLR). All calculations were done using RStudio (version 1.3.1093; https://www.rstudio.com/).

## Results

### fMRI

The activation maps of all participants revealed a moderate leftward lateralisation pattern of BOLD clusters (LI=0.36±0.263; Figure 5). Based on the LI of the fMRI language comprehension task in our ROIs, 906 participants were classed as left-hemisphere dominant (87%; LI= 0.43±0.174), 86 as bilateral (8%; LI=0.02±0.061), and the remaining 48 as right-hemisphere dominant (5%; LI=-0.38±0.259). The fMRI activation maps from these groups are shown in Figure 6. In both groups, strong neural activity is observed in brain areas associated with semantic processing i.e. anterior, and posterior temporal lobes as well as in the left inferior frontal gyrus (Jackson, 2021). In line with previous reports that have used similar story/narrative materials, also the ventral angular gyrus bordering the temporoparietal junction was engaged during language comprehension (Lerner et al, 2011; Branzi et al, 2020; Branzi et al., 2021; Humphreys & Lambon Ralph, 2015).

**Figure 5.**
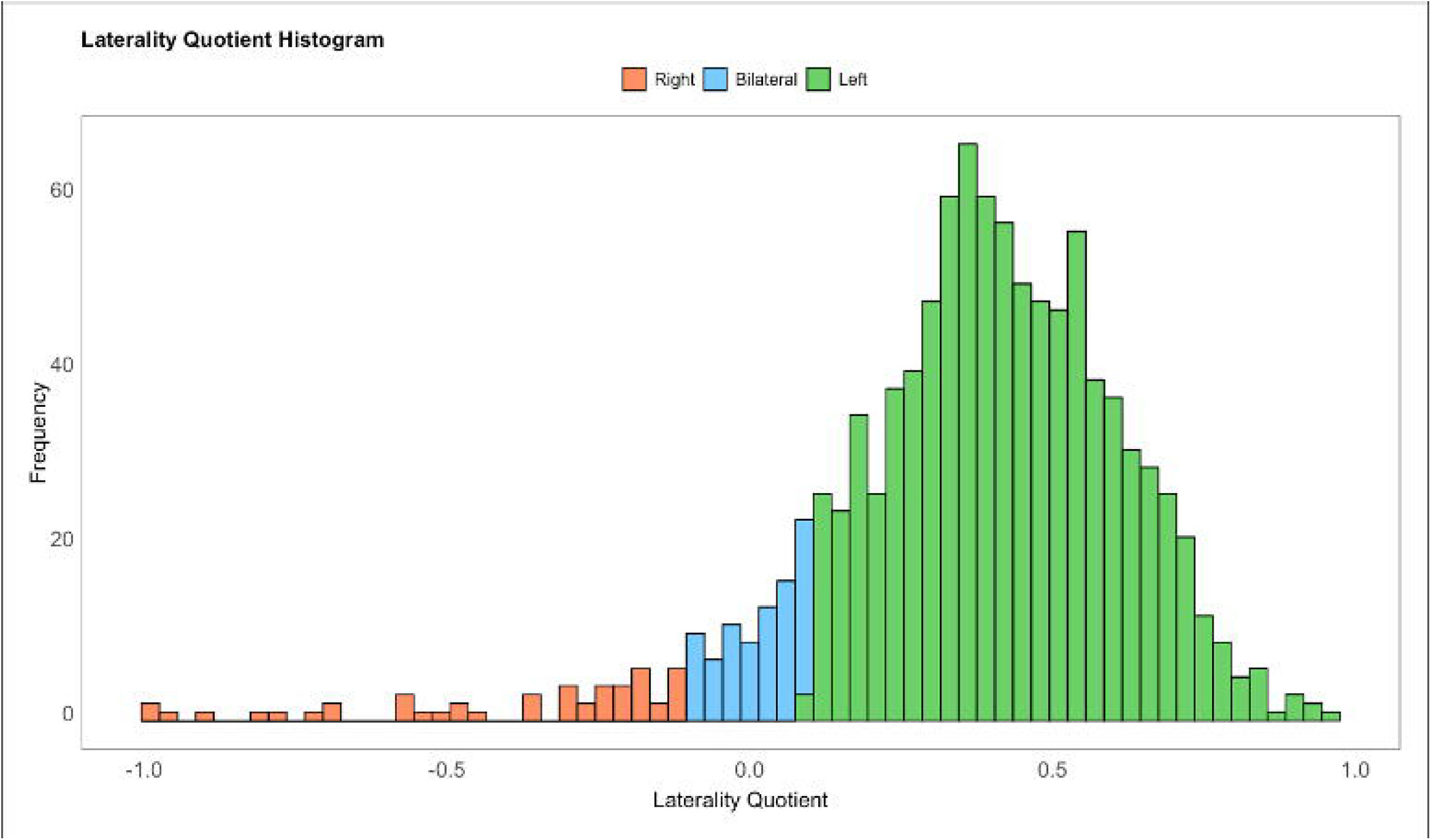
Histogram of laterality quotients in the whole sample (n=1040). Cut-off values were set at below −0.1 for right hemispheric dominance (red, n=48) and above 0.1 for left hemispheric dominance (green, n=906). Values in-between (i.e. −0.1 to 0.1) are considered as bilateral (i.e. no dominance, blue, n=86).

**Figure 6.**
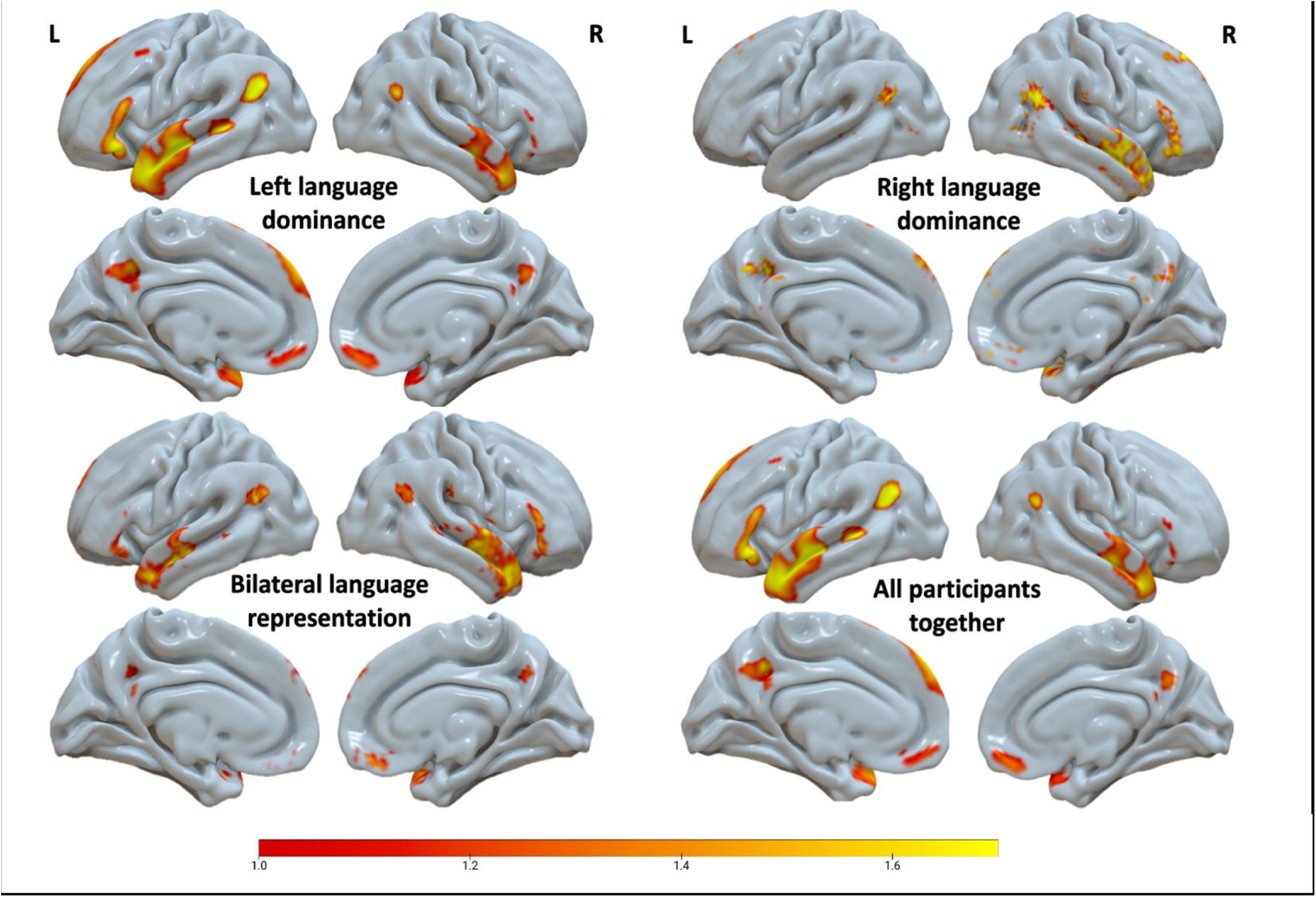
Cohen’s d maps of language reading comprehension task. The colour bar indicates Z scores; L – left hemisphere; R –right hemisphere. Brighter colours indicate stronger activation for the STORY > MATH contrast.

### Connectometry

We performed three connectometry analyses. First, in our study of individuals with bilateral and left hemisphere dominance (n=992), we identified 3,384 white matter tracts with a significant negative correlation with LQ. Specifically, reduced language dominance was associated with increased in QA in these tracts (p<0.05, FDR corrected) (Figure 7a). Notably, most tracts were commissural, accounting for 78% of the identified tracts. This included the forceps minor (51%), corpus callosum body (27%), forceps major (3%), and corpus callosum tapetum (1%). A smaller proportion of these tracts were situated in the fornix (20%) bundle bilaterally (Supplementary Fig. 1a). Conversely, we also observed a significant positive correlation with LQ in 339 tracts, demonstrating that a higher left hemisphere dominance was associated with increased QA in these tracts (p<0.05, FDR corrected) (Figure 7b). Among these tracts, 64% were localized within the left cingulum bundle, encompassing frontal parietal, frontal parahippocampal, and parolfactory areas. Furthermore, 12% were situated within the right cingulum bundle (aligning with regions observed in the left hemisphere), 13% in the right cerebellar tracts (including cerebellar peduncle and cerebellum), and 7% within the left arcuate fasciculus (Supplementary Fig. 1b). In the context of categorical group comparisons, our connectometry analysis revealed 412 tracts with increased QA in individuals with BLD compared to those with LLD (p<0.05, FDR corrected) (Supplementary Fig. 1c). The majority of significant tracts were commissural (52%), with the most prominent contributors being the forceps major (42%), corpus callosum tapetum (4%), corpus callosum body (4%), and forceps minor (2%). Other tracts were distributed bilaterally across the brainstem (34%), including corticospinal (25%), corticopontine (6%), and medial lemniscus (3%) bundles. A small proportion of streamlines were found in corticostriatal tract (4%) and right arcuate fasciculus (3%).

**Figure 7.**
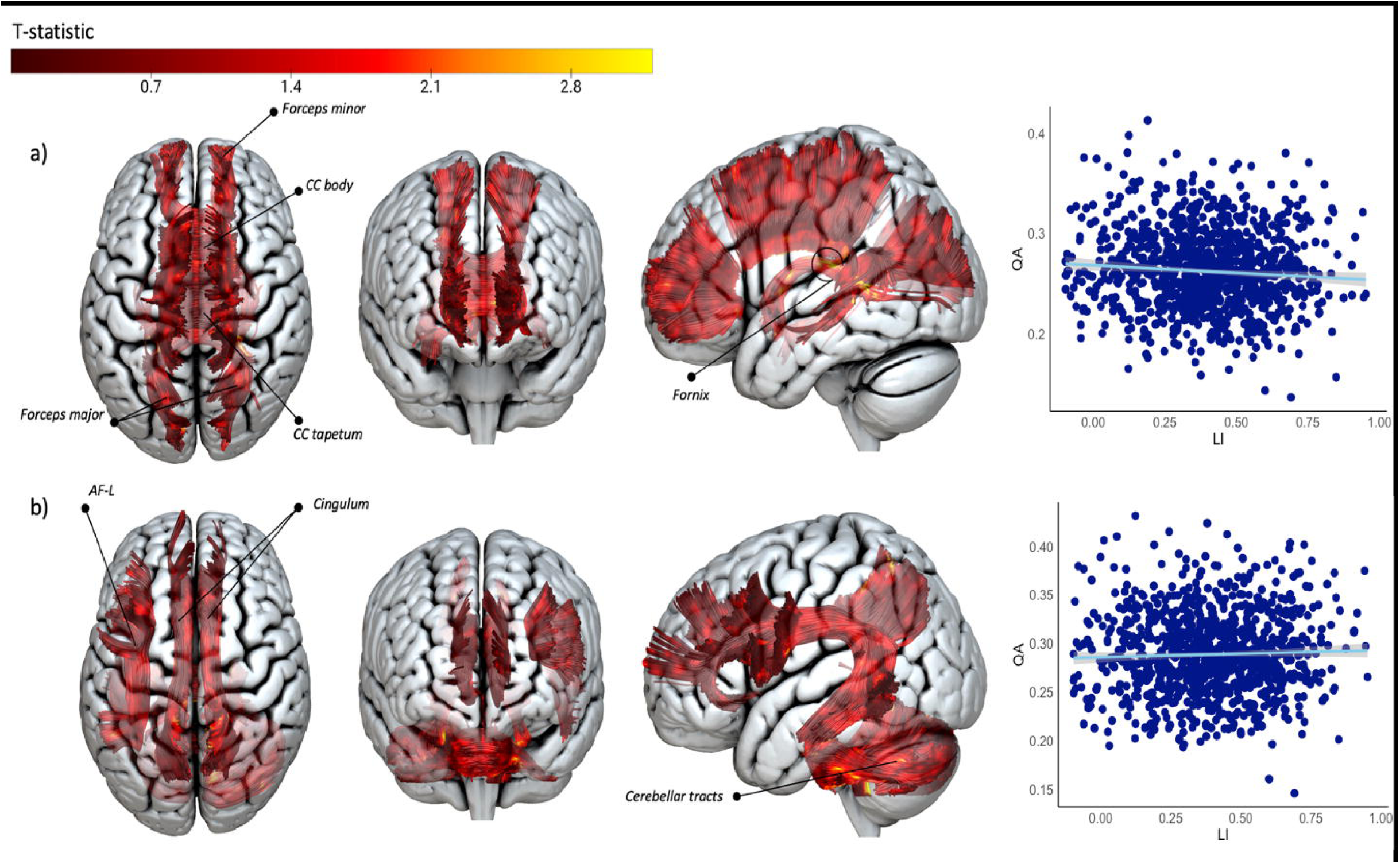
Connectometry results for the LLD and BLR groups (n=992). a) White matter bundles positively correlated with LQ (p<0.05, FDR corrected). b) White matter bundles negatively correlated with LQ (p<0.05, FDR corrected). Abbreviations: AF, arcuate e fasciculus; CC, corpus callosum; L, left hemisphere. In the visual representation, only tracts with more than 20 significant streamlines are shown as complete white matter bundles, with distinctive portions of the tract highlighted based on t-statistic colour to emphasise regions of significance. Scatter plots indicate the relationship between the mean QA in the corresponding tracts presented, and LQ. Each data point represents an individual subject.

In our second connectometry analysis, comprised of people with BLR and RLD (n=134), we identified 2,744 tracts positively correlated with LQ, indicative of a bilateral or near-zero laterality pattern (p<0.05, FDR corrected) (Figure 8). Most tracts were concentrated within the posterior commissural tracts (79%), which encompassed the forceps major (51%), corpus callosum tapetum (27%), and corpus callosum body (1%). The remaining tracts were distributed in the left inferior fronto-occipital fasciculus (6%), bilateral corticospinal tract (4%), right cerebellar tracts (4%) (including the middle cerebellar peduncle and cerebellum bundle), left superior corticostriatal tract (2%), and parietal corticopontine tract (1%) bilaterally (Supplementary Fig. 1a). When examining categorical group comparisons, we observed that 13 posterior commissural tracts, including the forceps major and tapetum, exhibited higher QA in bilateral individuals compared to those with left lateralization (p<0.05, FDR corrected) (Supplementary Fig. 1b).

**Figure 8.**
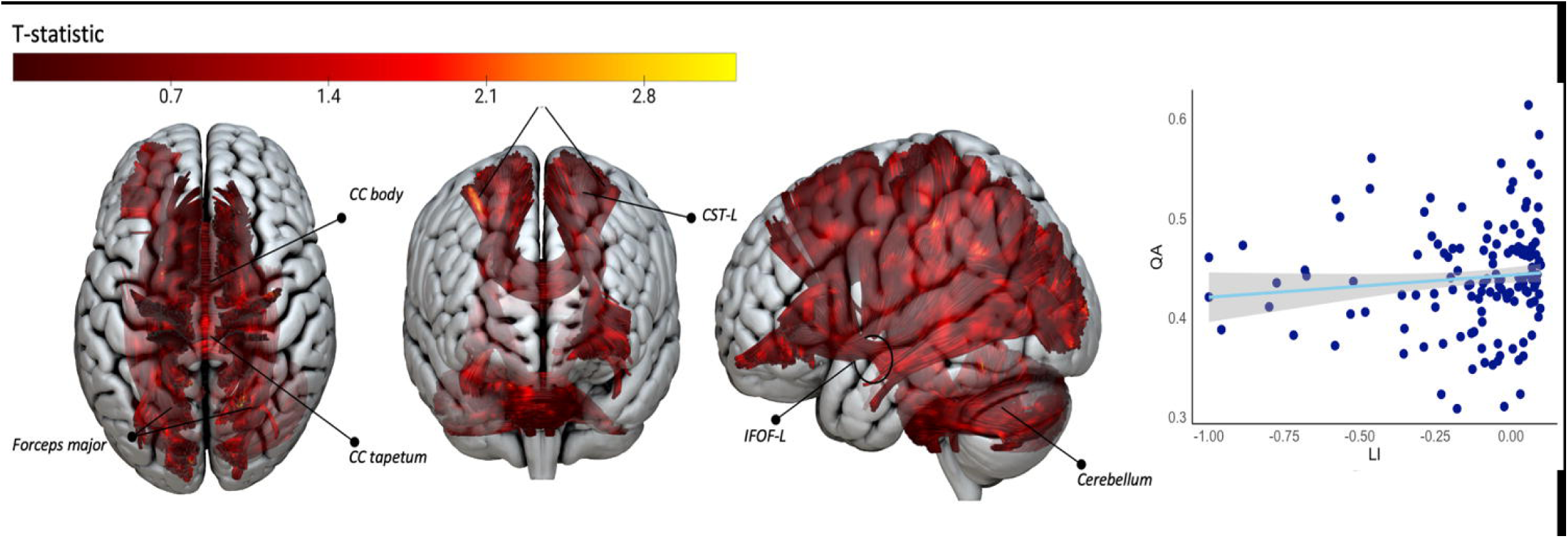
Connectometry results for the RLD and BLR groups (n=134), highlighting white matter bundles positively correlated with LQ (p<0.05, FDR corrected). Abbreviations: CC, corpus callosum; CPT, corticospinal tract; CST, corticostriatal tract; IFOF, inferior fronto-occipital fasciculus; L, left hemisphere. In the visual depiction, complete white matter bundles are showcased only for tracts with more than 20 significant streamlines, and distinctive portions of each tract are emphasised based on t-statistic colour to highlight regions of significance.

Finally, our third connectometry analysis, encompassing individuals with LLD and RLD (n=954), revealed no significant differences between those with LLD and RLD (p<0.05, FDR corrected).

### Shape analysis

Across various groups, a consistent pattern emerged with specific tracts exhibiting subtle associations with language lateralisation. Notably, the left dorsal SLF consistently showed positive correlations with LQ (indicative of LLD) with multiple shape metrics such as branch volume, mean length, curl, and elongation. Additionally, the right and left UF span and forceps minor elongation and curl also displayed consistent associations with left lateralisation, while a curl and a branch volume of right UF consistently indicated a more bilateral pattern of HLD (Supplementary Table 1). Categorical ANOVA analysis revealed subtle differences among various laterality groups in several right IFOF shape metrics, as well as in right AF mean length, left ventral SLF mean length and span, and right dorsal SLF volume (Supplementary Table 2). However, when corrected for false discovery rate (FDR), none of the aforementioned analyses yielded statistically significant results (Supplementary Table 1; Supplementary Table 2).

## Discussion

There were two main objectives of the present study. First, we investigated the relationship between degree of language lateralisation and QA within white matter tracts across the whole brain using connectometry. We found that reduced HLD was associated with increased QA, predominantly in the anterior callosal connections (forceps minor, body). The second objective was to determine whether interhemispheric asymmetries of white matter tract shape characteristics were associated with the degree of language lateralisation. After FDR correction, we found no significant correlation between length, span, curl, elongation, diameter, volume, and surface area of white matter tracts important for language and the degree of language lateralisation. We discuss the biological implications of these findings before addressing pertinent methodological issues.

### Biological implications

Our results indicate significantly increased QA of the corpus callosum (including forceps minor, body, tapetum, and forceps major) in people with BLR when compared to people with LLD. Similarly, bilateral individuals demonstrated higher QA in commissural tracts compared to those with right language laterality, but in contrast to previous group, this heightened involvement was primarily observed in the posterior streamlines (tapetum and forceps major). This observation aligns with prior research on brain tumour patients (Tantillo et al., 2016) and offers additional support to the hypothesis that individuals with enhanced reliance on both cerebral hemispheres for language processing might potentially possess more developed commissural fibres, this way compensating for the increased metabolic energy demands for information transfer and shortening the transmission times (Laughlin & Sejnowski, 2003). Notably, patients with BLR have been reported to have weaker memory, language skills, and lower IQ scores compared to lateralised individuals in some studies (Everts et al., 2009; Gleissner et al., 2003; Loring et al., 1999), whereas healthy individuals with BLD show the same level of cognitive performance as left-hemisphere dominant individuals in others (Knecht, 2001). The degree of lateralisation’s impact on cognitive measures has also been explored in the context of perisylvian white matter asymmetry. Catani et al. (2007) found that individuals with a more symmetric arcuate fasciculus in both hemispheres perform better in verbal memory tasks than those with asymmetric arcuate fasciculi.

Other imaging research has also suggested the importance of the corpus callosum for language lateralisation, although the link between structural measures of the corpus callosum and HLD is unclear. One study reported a greater fractional anisotropy of the whole corpus callosum in people with atypical language lateralisation (defined as RLD and BLR together) and, inconsistent with our results, greatest anisotropy in people with RLD (Häberling et al., 2011). Using a fixel-based approach, Verhelst and colleagues (2021) reported that people with RLD have greater fibre bundle cross-section of the right forceps minor compared to people with LLD (people with BLR were not investigated). The forceps minor is the group of anterior callosal connections, which pass through the genu of the corpus callosum and connect left and right prefrontal regions. These tracts are essential in integrating higher-order language and executive functioning (Hula et al., 2020; Mamiya et al., 2018; Sihvonen et al., 2021). Although commissural fibres have shown relationships with language lateralisation in the present and previous studies, there is inconsistency with respect to whom greater microstructural information inferred by the diffusion scalar metrics is reported. This may be due to methodological factors, such as the inclusion/exclusion of people with bilateral language representation and the type of language lateralisation assessed (see below).

Our study reported a significant association between the asymmetry in functional activation and QA in the fornix and corticostriatal tracts bilaterally, particularly in individuals exhibiting a LQ closer to zero, indicating a more bilateral hemispheric dominance. While there is limited existing literature exploring the direct relationship between LQ and diffusion measures in these tracts, a few studies have highlighted the involvement of them in language processing (Hula et al., 2020; Krishnan et al., 2016; Lee et al., 2013; Sihvonen et al., 2021). Conversely, we have identified a few streamlines (albeit a small number) that exhibit significant positive correlations between LQ and QA within a subgroup comprising individuals with LLD and BLR, but not within the group of individuals with RLD and BLR. These tracts include the left arcuate fasciculus, right cerebellar tracts, and bilateral cingulum, suggesting that an increase in QA in them may be associated with left-hemisphere language lateralization. Previous research has also suggested associations between left arcuate fasciculus asymmetry and LLD in several cohorts of healthy individuals (Ocklenburg et al., 2013; Propper et al., 2010), although the exact relationship between arcuate fasciculus asymmetry and HLD remains somewhat unclear (Gerrits et al., 2020; Verhelst et al., 2021). The observed right QA asymmetry in the cerebellar peduncle among individuals with LLD aligns with fMRI studies that have reported reversed functional activation patterns in the cerebellum (Starowicz-Filip et al., 2017; Stoodley and Schmahmann, 2009), however our study is the first to report such associations in diffusion-based metrics. As for the association between left-hemispheric language dominance and the cingulum, previous research has primarily been limited to one predictive study employing graph theory, which demonstrated that left hemisphere dominance can be predicted by a higher degree in the right posterior cingulate cortex (Silva and Citterio, 2017). Although this is the sole study to have identified associations between HLD and cingulum tracts, several other studies have indicated the involvement of this bundle in language processing (Forkel et al., 2022; Hula et al., 2020; Jung et al., 2019; Sihvonen et al., 2021). Further studies are needed better to understand these tracts’ importance for language lateralisation.

Several studies have investigated the relationship between characteristics of white matter and HLD, focusing on morphometric features (Josse et al., 2008; Karpychev et al., 2022; Westerhausen et al., 2006). However, no studies have combined tractography shape analysis with functional data in healthy individuals to examine the connection between geometric features of tracts and language lateralisation. A few subtle relationships were found in SLF, uncinate fasciculus, and forceps minor, although they served as poor predictors since none of them survived multiple comparisons correction. Therefore, we conclude that shape characteristics of language-relevant white matter tracts are not significantly associated with fMRI-based laterality measures. These results align with previous studies that found no relationship between language lateralisation and white matter features of arcuate fasciculus (Gerrits et al., 2022; Verhelst et al., 2021; Vernooij et al., 2007; Yazbek et al., 2021). This suggests that the well-documented left-hemisphere asymmetry of the arcuate fasciculus (Catani et al., 2007; Thiebaut de Schotten et al, 2011) may be detected using microstructural diffusion measures rather than shape features.

Notably, our shape analysis revealed no significant correlation between shape measures of the callosal tracts and language lateralisation, consistent with Westerhausen et al. (2006), who proposed that the microstructural organization, rather than the macrostructure, of the corpus callosum indicates functional language lateralisation. These findings differ from a recent study by Karpychev et al. (2022), which found that greater volumes of a posterior corpus callosum subregion were associated with a more substantial degree of language lateralisation (both RLD and LLD). Although subtle or no significant association was observed between long-range association fibres (including the inferior fronto-occipital fasciculus, superior longitudinal fasciculus, and uncinate fasciculus) and LQ in line with previous research (Delgado-Fernández et al., 2020; James et al., 2015; Silva & Citterio, 2017), a few earlier studies have reported conflicting results. For instance, Perlaki et al. (2013) identified a positive correlation between fractional anisotropy of the left superior longitudinal fasciculus and LI. However, their study solely examined the microstructural properties of the superior longitudinal fasciculus, making direct comparisons with our findings challenging. Additionally, Ocklenburg et al. (2014) reported a notable correlation between the volume of the right uncinate fasciculus and LI, although their study employed dichotic listening rather than a functional comprehension task.

### Methodological considerations

Although we employed traditional categorical classifications (left, right, bilateral) to assess language lateralisation, we also employed correlational techniques to investigate the association between LI and white matter characteristics. This approach mitigates the subjectivity concerns associated with the categorical grouping (Wegrzyn et al., 2019; Westerhausen et al., 2006). Even though, our connectometry findings remained consistent across both categorical and correlational analyses, continuous data analysis provided richer information for both BLR/LLD and BLR/RLD groups. For example, in the BLR and LLD group, correlational analysis revealed significant associations with left lateralization-related tracts, such as the left arcuate fasciculus, cingulum, and right cerebellar tracts, whereas categorical analysis found none. Similarly, in the BLR/RLD group, the correlational analysis offered more sensitivity by detecting corticostriatal, corticospinal, and corpus callosum body tracts that categorical analysis missed.

The number of individuals with non-left dominant language laterality, particularly right hemisphere dominant people, is small in the present study compared to previous studies (Chang et al., 2011). This disparity is primarily attributed to the task used to assess HLD. Specifically, perceptual tasks entailing semantic processing, such as the one adopted in this study, typically result in more bilateral fMRI activations (Binder et al., 2011; Metoki et al., 2022; Walenski et al., 2019). This might be due to several reasons. One possibility is that, as compared to the production of language, comprehension tasks rely less on the left-lateralised dorsal pathway, strongly associated with cognitive operations crucial for language production (e.g., retrieval and production of speech sounds) (Hickok & Poeppel 2007). Other non-mutually exclusive explanations include differences in the complexity of the linguistic stimuli (single words typically used in fluency tasks versus sentences typically used in comprehension tasks) and the demands that these tasks typically involve (see Peelle, 2012).

For example, the verbal stimuli used for the stories likely portray social concepts (e.g., theory of mind, intention, emotion, morality) and/or they may contain different amounts of metaphor, idiom, or implied meaning. All these aspects have been associated with the recruitment of fronto-temporal and parietal regions in the right hemisphere (Miller et al. 1997; Olson et al. 2007; Yang, 2014; Schmidt et al., 2007).

Finally, some difficulty in determining individuals with atypical lateralisation might be also due to the type of baseline task used in the present study. Indeed, previous studies that found left-lateralized semantic-related activations in fronto-temporal and parietal brain regions have utilized as baseline reversed/rotated speech or pseudowords which share various linguistic representations with words, except for semantics (see Peelle, 2012).

Be that as it may, the fact we employed a language comprehension task might explain why only a small subset of participants demonstrated strong leftward lateralization, while the majority displayed mild lateralization. A recent systematic review comparing different language tasks has highlighted that language production tasks may be more robust in accurately assessing language laterality than language comprehension tasks (Bradshaw et al., 2017). For instance, there is evidence that between 6 and 24 years of age, there is an increase in frontal asymmetry during tasks involving the articulation of words. However, this asymmetry is not present during story listening. This suggests partly different maturational mechanisms between language comprehension and production (Lidzba et al., 2011; Berl et al., 2010). Future studies will have to use tasks that typically generate strongly left lateralised neural responses such as verbal fluency tasks to corroborate our findings.

Our methodology offers several practical benefits. Firstly, despite the limited proportion of right hemisphere dominant individuals identified through the functional language task, the sizeable sample size (N= 1040) employed in this study is the largest ever utilized in such investigations, thus potentially decreasing the likelihood of false positives. Furthermore, we employed two complementary methodologies, namely connectometry and tractography, to delineate the precise white matter characteristics associated with HLD. In doing so, we have provided new insights into the anatomical basis of language lateralisation insomuch that white matter tract macrostructural geometrical features (shape analysis) are unrelated to HLD at the group-level.

## Conclusion

The findings of our study suggest that measures of diffusion-based microstructural architecture, rather than the geometric characteristics of reconstructed white matter tracts, are linked to language lateralisation. Specifically, individuals who exhibit a greater reliance on both cerebral hemispheres for language processing may possess more highly developed commissural fibres, thereby promoting more efficient interhemispheric communication.

## Supporting information

Supplemental data description

## Funding/Acknowledgements

The research was supported by a BBSRC Studentship and UKRI Future Leaders Fellowship (MR/T04294X/1).

