## Supplementary figures and images for "The relationship between white matter architecture and language lateralisation in the healthy brain"

### Supplemental Fig. 1

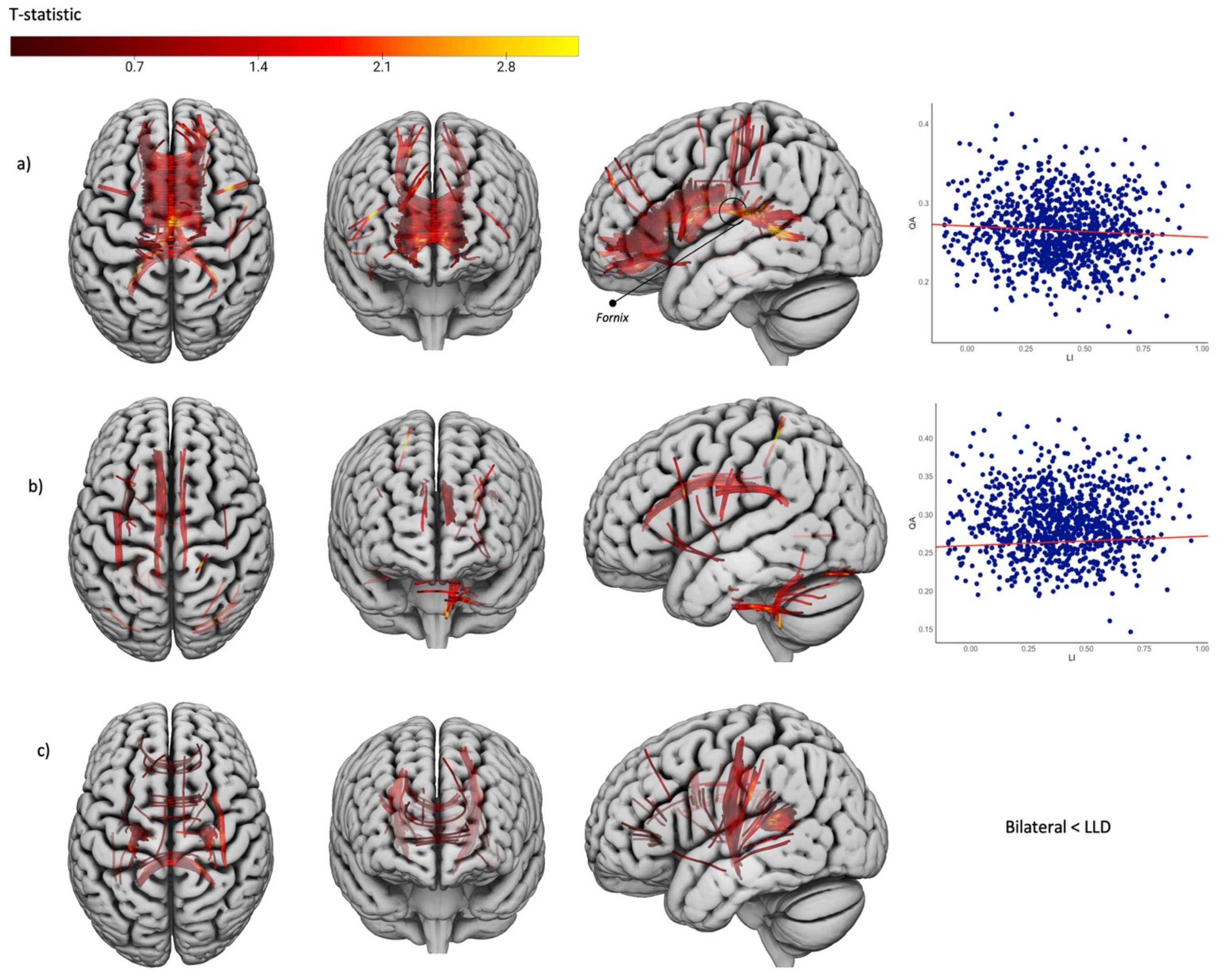

### Supplemental Fig. 2

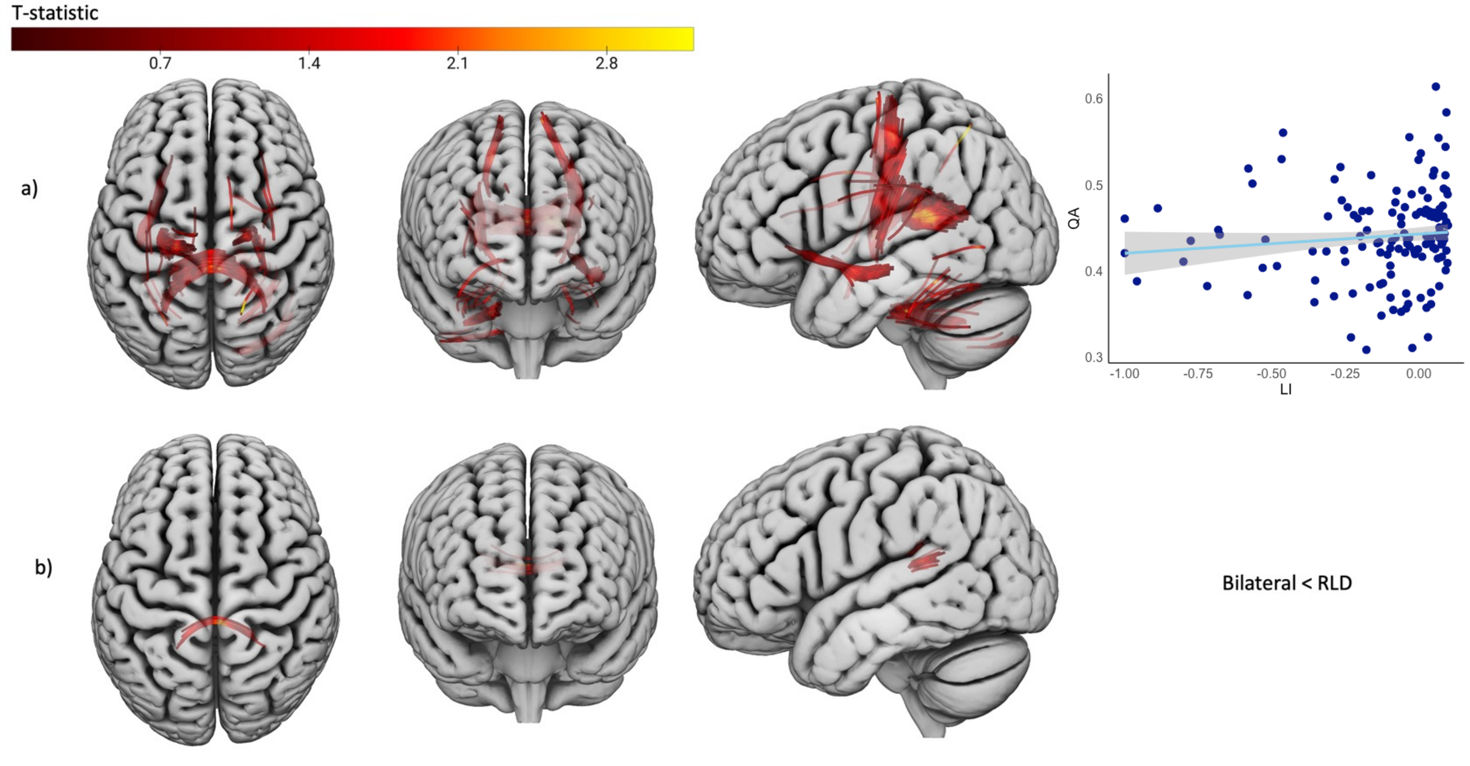
