## Supplemental Table 1 for "The relationship between white matter architecture and language lateralisation in the healthy brain"

| Tract | Measure | P Value | Associated with? | Group | FDR |
|---|---|---|---|---|---|
| Forceps minor | diameter(mm) | 0.005 | BLR | LLD/BLR | 0.25 |
| Forceps minor | elongation | 0.01 | LLD | LLD/BLR | 0.372 |
| Forceps minor | curl | 0.029 | LLD | LLD/BLR | 0.472 |
| Forceps minor | volume(mm^3) | 0.04 | BLR | LLD/BLR | 0.472 |
| Forceps minor | diameter(mm) | 0.037 | RLD | LLD/RLD | 0.741 |
| Forceps minor | volume(mm^3) | 0.039 | RLD | LLD/RLD | 0.741 |
| IFOF L | elongation | 0.037 | LLD | LLD/BLR | 0.472 |
| IFOF L | elongation | 0.018 | LLD | LLD/RLD | 0.741 |
| IFOF R | volume(mm^3) | 0.009 | RLD | RLD/BLR | 0.656 |
| IFOF R | total surface area(mm^2) | 0.017 | RLD | RLD/BLR | 0.656 |
| IFOF R | trunk volume(mm^3) | 0.018 | RLD | RLD/BLR | 0.656 |
| IFOF R | diameter(mm) | 0.021 | RLD | RLD/BLR | 0.656 |
| SLF 1 L | branch volume(mm^3) | 0.001 | LLD | LLD/BLR | 0.078 |
| SLF 1 L | mean length(mm) | 0.016 | LLD | LLD/BLR | 0.472 |
| SLF 1 L | curl | 0.034 | LLD | LLD/BLR | 0.472 |
| SLF 1 L | elongation | 0.043 | LLD | LLD/BLR | 0.472 |
| SLF 1 L | branch volume(mm^3) | 0.014 | RLD | RLD/BLR | 0.656 |
| SLF 1 R | volume(mm^3) | 0.037 | RLD | LLD/RLD | 0.741 |
| SLF 2 L | volume(mm^3) | 0.043 | BLR | LLD/BLR | 0.472 |
| SLF 2 L | total surface area(mm^2) | 0.026 | RLD | RLD/BLR | 0.701 |
| SLF 2 L | total surface area(mm^2) | 0.028 | RLD | LLD/RLD | 0.741 |
| SLF 2 L | volume(mm^3) | 0.037 | RLD | LLD/RLD | 0.741 |
| SLF 2 R | trunk volume(mm^3) | 0.042 | BLR | RLD/BLR | 0.94 |
| SLF 3 L | volume(mm^3) | 0.024 | LLD | LLD/BLR | 0.472 |
| SLF 3 L | span(mm) | 0.026 | LLD | LLD/BLR | 0.472 |
| SLF 3 L | curl | 0.044 | BLR | LLD/BLR | 0.472 |
| SLF 3 R | curl | 0.035 | BLR | LLD/BLR | 0.472 |
| SLF 3 R | total surface area(mm^2) | 0.036 | LLD | LLD/RLD | 0.741 |
| UF L | span(mm) | 0.02 | LLD | LLD/BLR | 0.472 |
| UF R | span(mm) | 0.001 | LLD | LLD/BLR | 0.078 |
| UF R | curl | 0.001 | BLR | LLD/BLR | 0.078 |
| UF R | branch volume(mm^3) | 0.016 | BLR | RLD/BLR | 0.656 |
| UF R | curl | 0.009 | RLD | LLD/RLD | 0.741 |
| UF R | span(mm) | 0.036 | LLD | LLD/RLD | 0.741 |
