## Supplemental Table 2 for "The relationship between white matter architecture and language lateralisation in the healthy brain"

| Tract | Measure | F Value | P Value | FDR |
|---|---|---|---|---|
| AF R | mean length(mm) | 5.159 | 0.006 | 0.26 |
| IFOF R | trunk volume(mm^3) | 5.816 | 0.003 | 0.26 |
| IFOF R | volume(mm^3) | 5.611 | 0.004 | 0.26 |
| IFOF R | total surface area(mm^2) | 5.128 | 0.006 | 0.26 |
| IFOF R | diameter(mm) | 3.907 | 0.02 | 0.436 |
| IFOF R | mean length(mm) | 3.142 | 0.044 | 0.747 |
| SLF1 R | volume(mm^3) | 4.214 | 0.015 | 0.368 |
| SLF3 L | mean length(mm) | 4.87 | 0.008 | 0.269 |
| SLF3 L | span(mm) | 4.38 | 0.013 | 0.364 |
