## Supplemental data description for "The relationship between white matter architecture and language lateralisation in the healthy brain"

**Supplementary Figure 1.** Raw connectometry results for the LLD and BLR groups (n=992). a) Tract sections positively correlated with LQ (p<0.05, FDR corrected). b) Tract sections negatively correlated with LQ (p<0.05, FDR corrected). c) Tract sections associated with bilateral individuals compared to LLD (p<0.05, FDR corrected).

**Supplementary Figure 2.** Raw connectometry results for the RLD and BLR groups (n=134). a) Tract sections positively correlated with LQ (p<0.05, FDR corrected). b) Tract sections associated with bilateral individuals compared to RLD (p<0.05, FDR corrected).

**Supplementary Table 1.** Summary of regression analysis results with uncorrected p-values across all groups. IFOF, inferior fronto-occipital fasciculus; SLF, superior longitudinal fasciculus; UF, uncinate fasciculus; L, left hemisphere; R, right hemisphere.

**Supplementary Table 2.** Summary of significant uncorrected ANOVA results. AF, arcuate fasciculus; IFOF, inferior fronto-occipital fasciculus; SLF, superior longitudinal fasciculus; L, left hemisphere; R, right hemisphere.
